# Antagonism Between Brain Regions Relevant for Cognitive Control and Emotional Memory Facilitates the Generation of Humorous Ideas

**DOI:** 10.1101/2020.06.03.131417

**Authors:** Florian Bitsch, Philipp Berger, Andreas Fink, Arne Nagels, Benjamin Straube, Irina Falkenberg

**Affiliations:** Department of Psychiatry and Psychotherapy, Philipps-University Marburg, Rudolf-Bultmann Str. 8, 35039 Marburg, Germany; Center for Mind, Brain and Behavior – CMBB, Hans-Meerwein-Straße 6, 35032, Marburg, Germany; Department of Neuropsychology, Max Planck Institute for Human Cognitive and Brain Sciences, Stephanstraße 1a, 04103 Leipzig, Germany; Institute of Psychology, University of Graz, BioTechMed, Universitätsplatz 2, 8010, Graz, Austria; Department of English and Linguistics, Johannes Gutenberg-University Mainz, Jakob-Welder-Weg 18, 55128 Mainz, Germany

**Keywords:** Humor Generation, Creativity, Cognitive Control, fMRI, AU Task, Stroop Task, Inhibition

## Abstract

The ability to generate humor gives rise to positive emotions and thus facilitate the successful resolution of adversity. Although there is consensus that inhibitory processes might be related to broaden the way of thinking, the neural underpinnings of these mechanisms are largely unknown. Here, we use functional Magnetic Resonance Imaging, a humorous alternative uses task and a stroop task, to investigate the brain mechanisms underlying the emergence of humorous ideas in 24 subjects. Neuroimaging results indicate that greater cognitive control abilities are associated with increased activation in the amygdala, the hippocampus and the superior and medial frontal gyrus during the generation of humorous ideas. Examining the neural mechanisms more closely shows that the downregulation of frontal brain regions is associated with an upregulation in the amygdala, which is concurrently linked with an increased number of humorous ideas and enhanced amygdala responses during the task. Our data therefore suggests that a neural antagonism previously related to the emergence and regulation of negative affective responses, is linked with the generation of emotionally positive ideas and may represent an important neural pathway supporting mental health.

## Introduction

Creative thinking can be considered a central building block of human achievements since it promotes the emergence of novel concepts by re-organizing established principles. One important aspect of creative thinking is the generation of humorous ideas (Ruch & Heintz, 2019), which can be regarded as an emotionally positively laden component of divergent thinking (Benedek & Fink, 2019). Although the ability is deeply grounded in human social life, facilitating social relationships (Berger et al., 2018a) and promoting mental health (Samson et al., 2014; Falkenberg et al., 2013), its neural mechanisms are only beginning to be understood (e.g., Perchtold-Stefan et al., 2020). Studies examining the cognitive and neural mechanisms of divergent thinking have pointed out that cognitive control mechanisms can facilitate the mining of abstract ideas (Jung et al., 2013). However, it is currently unclear, whether high (e.g. Edl et al., 2014) or low inhibitory mechanisms (Radel et al., 2015) enable the creative process (Benedek & Fink, 2019) and how these are represented in the brain. To shed light on the cognitive and neural mechanisms underlying the ability to generate humorous ideas, we use functional neuroimaging to examine humor generation and how it may be facilitated by inhibitory capacities.

In the context of divergent thinking it has been suggested that the generation of creative ideas is closely interweaved with basic cognitive processes, such as cognitive control and executive functions (Jung et al., 2013; Zabelina & Robinson, 2010). More specifically, it has been assumed that during the generation of creative ideas, an active inhibition of content-related semantic mental representations facilitates a deeper search process to combine remote and therefore unusual knowledge structures within an associative semantic network (Mednick, 1962; Mohr et al., 2001). This line of thinking received empirical support from studies showing that higher individual cognitive control capacities are associated with the formation of novel and abstract concepts (Benedek et al., 2014). Along these lines, some studies demonstrated that more creative individuals can be characterized by more flexible cognitive control, as indicated by greater trial-to-trial cognitive control modulation (Zabelina & Robinson, 2010; see also Zabelina & Ganis, 2018). Furthermore, in a group of creative design students, an increased production of creative ideas was related with enhanced cognitive control abilities (Edl et al., 2014). Interestingly, however, although broad consensus exists that cognitive control processes are linked with the re-combination of established concepts to generate novel ideas, their specific contribution to the process of divergent thinking remains controversial (Benedek & Fink, 2019). Another line of research suggests that a reduction of cognitive control per se or a decoupling of inhibitory mechanisms is linked with a creative mind. This assumption was driven by findings of an overrepresentation of creative individuals in specific mental disorders in which dis-inhibition is a critical psychopathological facet, such as schizophrenia and bipolar disorders (Acar et al., 2017; Carson, 2011; Dietrich, 2014; Fink et al., 2014; Kyaga et al., 2013). To simulate such a disinhibition effect, healthy subjects received a mild alcohol intoxication which led to a reduction in cognitive control capacities, but an increase in creative processes (Benedek et al., 2017). Along these lines, experimentally induced reduction of inhibitory cognitive control capabilities by exhaustive practice of related cognitive tasks, led to a higher originality and frequency of abstract ideas (Radel et al., 2015). While these findings suggest close links between idea generation processes and inhibitory mechanisms during the mining of abstract ideas, brain imaging techniques can provide additional insights into their interrelationship, thereby facilitating a more comprehensive understanding of the manifold neurocognitive mechanisms implicated in this complex mental ability domain.

During creative cognition, neuroimaging findings reveal activity patterns in a widespread neural network involving brain regions related with cognitive control mechanisms such as the superior frontal gyrus (Hu et al., 2016), the inferior frontal gyrus (IFG, Beaty et al., 2014; Flaherty, 2005), the dorsolateral prefrontal cortex (dlPFC, Beaty et al., 2015; Flaherty, 2005) and brain regions associated with default mode processes such as the bilateral inferior parietal lobule (IPL), the posterior cingulate cortex (PCC), the medial prefrontal cortex (mPFC) and areas in the medial temporal lobe (MTL) such as the amygdala and the hippocampus, which are assumed to be critical for spontaneous and broadened thinking (Bitsch et al., 2018a; Bitsch et al., 2018b; Buckner et al., 2008). Humor production abilities have been found to have much in common with other forms of verbal creativity (Nusbaum et al., 2017; Perchtold-Stefan et al., 2020). Accordingly, similar functional networks of brain regions are assumed in association with the production of humorous ideas (e.g. amygdala and hippocampus), and with humor comprehension processes (e.g. IFG and IPL; Samson et al., 2009; Vrticka et al., 2013). Among these brain regions, the amygdala might have a central role for mining abstract concepts, given its implication for the acquisition, storage and modulation of emotional memory information (Ehrlich et al., 2009). Furthermore, the region is a central area in the processing of humor (Berger et al., 2018b; Iidaka, 2017). The degree to which inhibitory processes influence the regions’ functions during the creative process (Flaherty, 2005) is yet unknown. Previous research however suggests that the amygdala function is balanced by inhibition and disinhibition processes resulting in enhanced or reduced formation of emotions and cognition such as anxiety and fear (Babaev, et al., 2018; Ehrlich et al., 2009), a process particularly modulated by brain regions in the frontal cortex (Gee et al., 2013; Gold et al., 2014).

Thus, a better understanding of whether inhibitory mechanisms modulate the mining of humorous ideas would shed light on the relationship between inhibitory mechanisms and memory-informed spontaneous cognition (Benedek & Fink, 2019) and extent our understanding of the mechanisms underlying humor generation. Here, we use a modified version of the alternative uses (AU) task (e.g. Fink et al., 2011) as a well-established paradigm to assess the emergence of funny ideas. During the fMRI-task participants generated ideas for either funny or typical uses of everyday objects, while speech production was audiorecorded on-line. To examine whether inhibitory mechanisms modulate the mining of humorous ideas, we assessed each participant’s cognitive control capacities with a color naming stroop task outside the MRI and examined their potential effect on the idea generation process. We hypothesize that brain regions relevant for processing humor, such as the prefrontal cortex, the amygdala and the hippocampus, are implicated in the generation of funny (vs. typical) ideas. Furthermore, we expected that these brain regions and particularly prefrontal brain regions are modulated by inhibitory mechanisms and facilitate the generation of humorous ideas. Last, based on studies showing the high relevance of the frontal gyrus (e.g. Chan et al., 2013) and the amygdala (e.g. Berger et al., 2018b), we assumed that functional connectivity between the both regions will facilitate the flourishing of humorous ideas.

## Materials and methods

### 2.1 Participants

24 participants (12 females) between 23 and 56 years of age (*M±STD*=32.71±9.66) participated in the study. The sample was recruited via advertisements and e-mail distributors from (and to represent) the general population. All participants had normal or corrected-to-normal vision, reported no history of neurological or psychiatric disorders (which was excluded by a structured clinical interview for DSM-IV) and were right-handed according to Edinburgh Handedness Inventory (Oldfield, 1971). The study was approved by the local Ethics Committee at Philipps-University Marburg. All participants gave written informed consent prior to the experiment and were reimbursed for the time they spent at the study site.

### 2.2 Experimental task

Participants performed an adapted AU task, where they were presented with 30 names of everyday objects written in either green (funny (FUN) condition, 15 items) or red (typical (TYP) condition, 15 items) font color for the duration of 3 seconds, followed by a speech bubble for the duration of 12 seconds. Participants were instructed to generate a high quantity and quality of humorous ideas or typical uses of the presented items, depending on the condition, as soon as the speech bubble occurred (see Figure 1). The participants verbally reported the ideas that spontaneously came to their mind e. g. balloon: “I put it under my shirt that it seems that I am overweighted or pregnant” (FUN condition). Or in the TYP condition: e.g. soap: “I use it to wash myself’.

**Figure 1.**
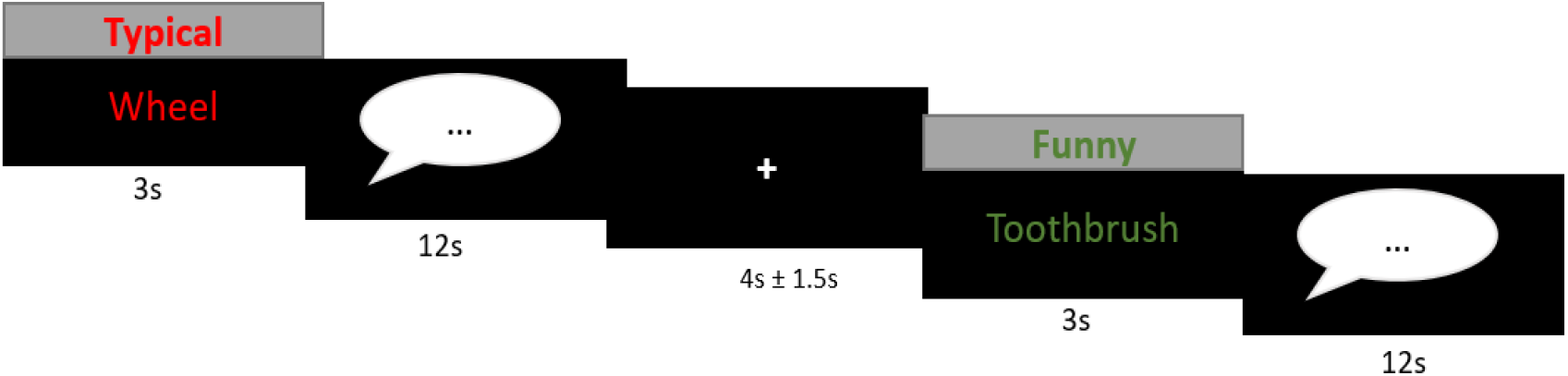
Experimental paradigm. During the task participants had to find either funny or typical usages for everyday objects (15 objects per condition). First participants were presented with the condition and word, afterwards they had to think aloud their idea. The ideas were recorded with a fMRI-suitable microphone and on-line transcripted.

Verbal answers were audio-taped during the scanning procedure and verbal output during the speech phase was transcripted using FOLKER (Schmidt & Schütte, 2010). Afterwards, the amount of i) generated words and ii) ideas (fluency score) was assessed. For each item, the amount of generated ideas was rated as follows: e.g. for the item “picture frame” a participant responded: “Carving the picture-frame by oneself, put oneself into the picture frame”. This answer was counted as two produced ideas for this participant and item.

### 2.3 Individual Cognitive Control Abilities

To assess the participant’s ability to inhibit cognitive interference of impeding stimuli-features, we applied the Stroop Color and Word Test, outside and following the MRI. The task was programmed and run within the Psychtoolbox implemented in Matlab (http://psychtoolbox.org/). The task included 4 words and colors: blue, green, red, yellow and the participants had to respond within 1.3 seconds to the color displayed (with one of four keys). The stroop interference score (Stroop, 1935), which is the ratio between the correct reaction times in the congruent condition (word and color identical) vs. the incongruent condition (word and color differently), was subsequently calculated and used a as an indicator for each participant’s inhibitory performance.

### 2.4 Individual Differences in Emotion Regulation and Coping

To examine the participants’ differences in regulating emotional experiences and stressful life-events, we assessed the emotion regulation questionnaire (ERQ, Gross & John, 2003) following the fMRI-session. The ERQ is a 10-items self-report which measures two behavioral tendencies how individuals make use of two different emotion regulation process either reappraisal (6-items) or suppression (4-items).

### 2.5 Imaging procedure

#### 2.5.1 MRI data acquisition

All images were acquired using a Siemens 3-Tesla Trio with a 12-channel head matrix receive coil. Functional images were acquired using a T2* weighted single shot echo planar imaging (EPI) sequence (parallel imaging factor of 2 (GRAPPA), TE=30ms, TR=2000ms, flip angle 90°, slice thickness 3.6 mm, matrix 64×64, in-plane resolution 3×3 mm^2^, bandwidth 2232 Hz/pixel, EPI factor of 64 and an echo spacing of 0.51 ms). Data from 33 transversal slices oriented to the AC–PC line were gathered in descending order. Additionally, a whole head T1 weighted data set was acquired with a 3D MP-Rage sequence (parallel imaging factor of 2 (GRAPPA), TE=2.26ms, TR=1900ms, flip angle 9°, 1mm isometric resolution, 176 sagittal slices, 256×256 matrix).

#### 2.5.2 fMRI data preprocessing

Functional data preprocessing and analysis were performed using SPM12 (http://www.fil.ion.ucl.ac.uk/spm) implemented in MATLAB 13a (MathWorks, MA). The first five volumes of each functional run were discarded from the analysis to account for T1 equilibration effects. Functional data were realigned and unwarped, corrected for slice timing, spatially normalized onto a common brain space (Montreal Neurological Institute, MNI) and spatially smoothed using a Gaussian filter with an 8 mm full-width half maximum (FWHM) Gaussian kernel. In order to improve fMRI-data quality, an automated quality assurance protocol (Artifact (ART) Detection toolbox, https://www.nitrc.org/projects/artifact_detect/), based on head motion (> 2mm) or aberrant signal intensity (global-signal z-value exceeded a threshold of 9) was run and the resulting parameters were subsequently included in the subsequent fMRI analysis.

#### 2.5.3 fMRI data analysis

For statistical analysis we set up a general linear model (GLM) in SPM12 on the subject level, with 4 regressors comprising the onsets of the four task events (word: funny and typical (3 sec), speech: funny and typical (12 sec), Fig. 1). The number of words spoken were included as parametric modulator in the speech condition to control for speech related activity. Subsequently, the data were high-pass filtered with a cut-off frequency of 128s. To control for movement related activity, the realignment parameters and art parameters (see methods section 2.5.2) were included in the first-level model. Contrasts of interests were the funny > typical condition (speech and word phase), which were calculated during the first-level analysis. Afterwards the related contrast images were included in a second level model (one sample t-test). To correct for multiple voxel comparisons an extent threshold was used via Monte–Carlo simulation (Slotnick et al., 2003). The whole-brain activation was simulated assuming a type I error voxel activation of p=.001, this revealed a cluster extent of 47 contiguous resampled voxels as sufficient to correct for multiple comparisons at p<.05.

#### 2.5.4 Functional Analysis of Covariance

To investigate the relationship between inhibitory skills and the creative idea generation process, we additionally conducted covariate analyses by including the individual stroop scores (see the methods section 2.3) as a covariate of interest in a separate second level GLM. One participant has not conducted the stroop task and was therefore excluded from the covariate analysis.

#### 2.5.5 Functional Connectivity

We assessed task-related connectivity changes with the generalized form of contextdependent psychophysiological interaction (gPPI, McLaren et al., 2012). The gPPI approach is appropriate when more than two conditions are modeled in the design matrix, as higher sensitivity and specificity due to a better model fit compared to the standard form of PPI analysis suggests (McLaren et al., 2012). For the connectivity analysis, a functionally defined region of interest (ROI) of the superior/medial frontal gyrus from the analysis of covariance (please see: results section 3.2.3) was used as seed region. The region’s eigenvariate was extracted and for each condition a PPI-regressor was built and included in a first-level model. This first-level model consisted of the regressors from the functional activity analysis (described above) in addition to the seed-region’s time course. Contrast maps for FUN>TYP were calculated subsequently for each participant and submitted to a second-level random effects analysis (one sample *t*-test) including the stroop score as a covariate, analogous to the functional activity analysis.

#### 2.5.6 Correlation Analysis

In order to examine a potential relationship between functional activity and functional connectivity measures with the idea generation processes (quantity of funny ideas), we extracted the beta estimates from the entire functionally defined cluster of the amygdala and the superior/medial frontal gyrus and conducted subsequent correlation analysis by skipped correlations (Pernet et al., 2013; Wilcox, 2004) using the Matlab based robust correlation toolbox (http://sourceforge.net/projects/robustcorrtool/), which detects and disregards outliers depending on the data structure (Pernet et al., 2013).

## Results

### 3.1 Behavioral analysis

On average the participants generated *M*±STD=1.82±0.73 ideas per item in the funny condition and *M*±STD=2.27± 0.79 ideas per item in the typical condition. The higher amount of generated ideas in the typical vs the funny condition, *t*(23)= 2.52, p= 0.02, suggests that participants produced ideas more easily in the typical condition.

### 3.2 fMRI analysis

#### 3.2.1 Functional Activity Analysis: FUN>TYP

The whole-brain analysis of funny versus typical idea generation (FUN>TYP) revealed activation of a network spanning cortical and subcortical as well as cerebellar areas, including the amygdala/hippocampus complex, superior frontal and medial gyrus, the anterior cingulate cortex (ACC), the inferior frontal gyrus (IFG) and the temporal pole (Fig. 2, Tab.1).

**Figure 2.**
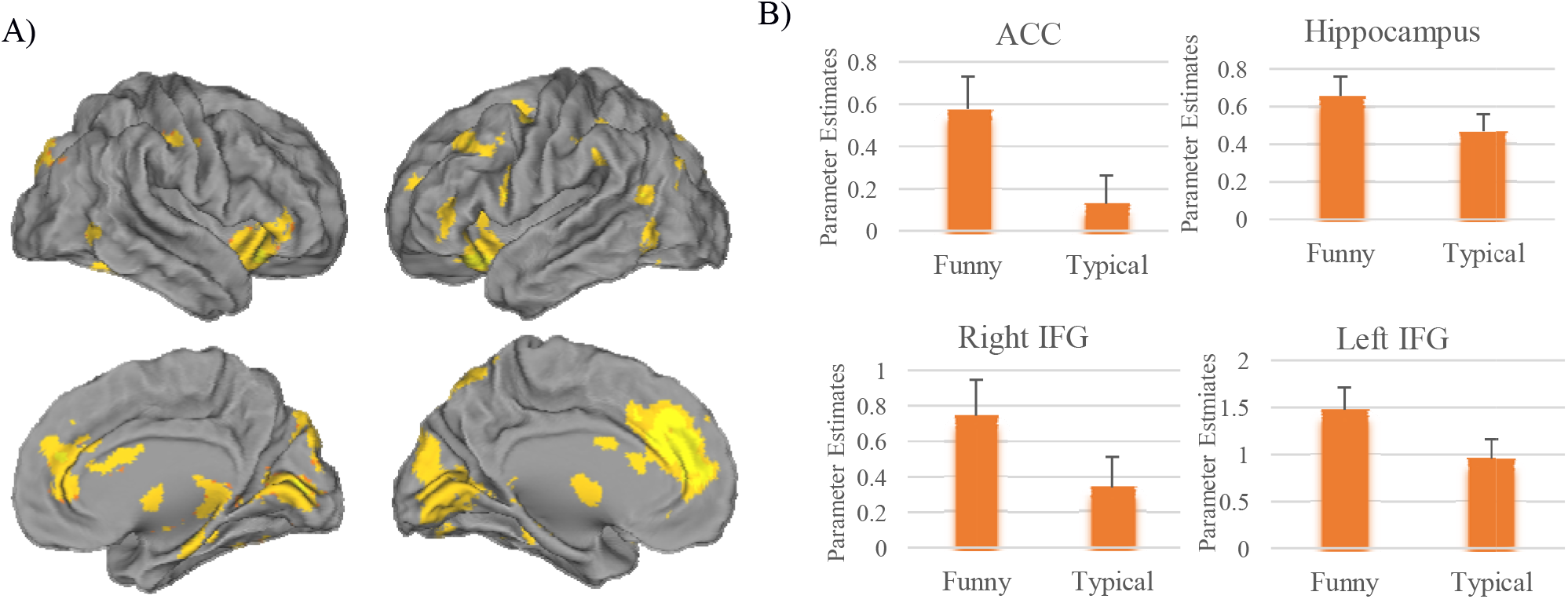
Functional Activity Analysis: A) The functional activity analysis of the generating FUN>TYP ideas contrast shows activity in a set of fronto-temporal brain regions, such as B) the ACC, the bilateral IFG and the left hippocampus. The error bars represent SEM.

**Table 1.**
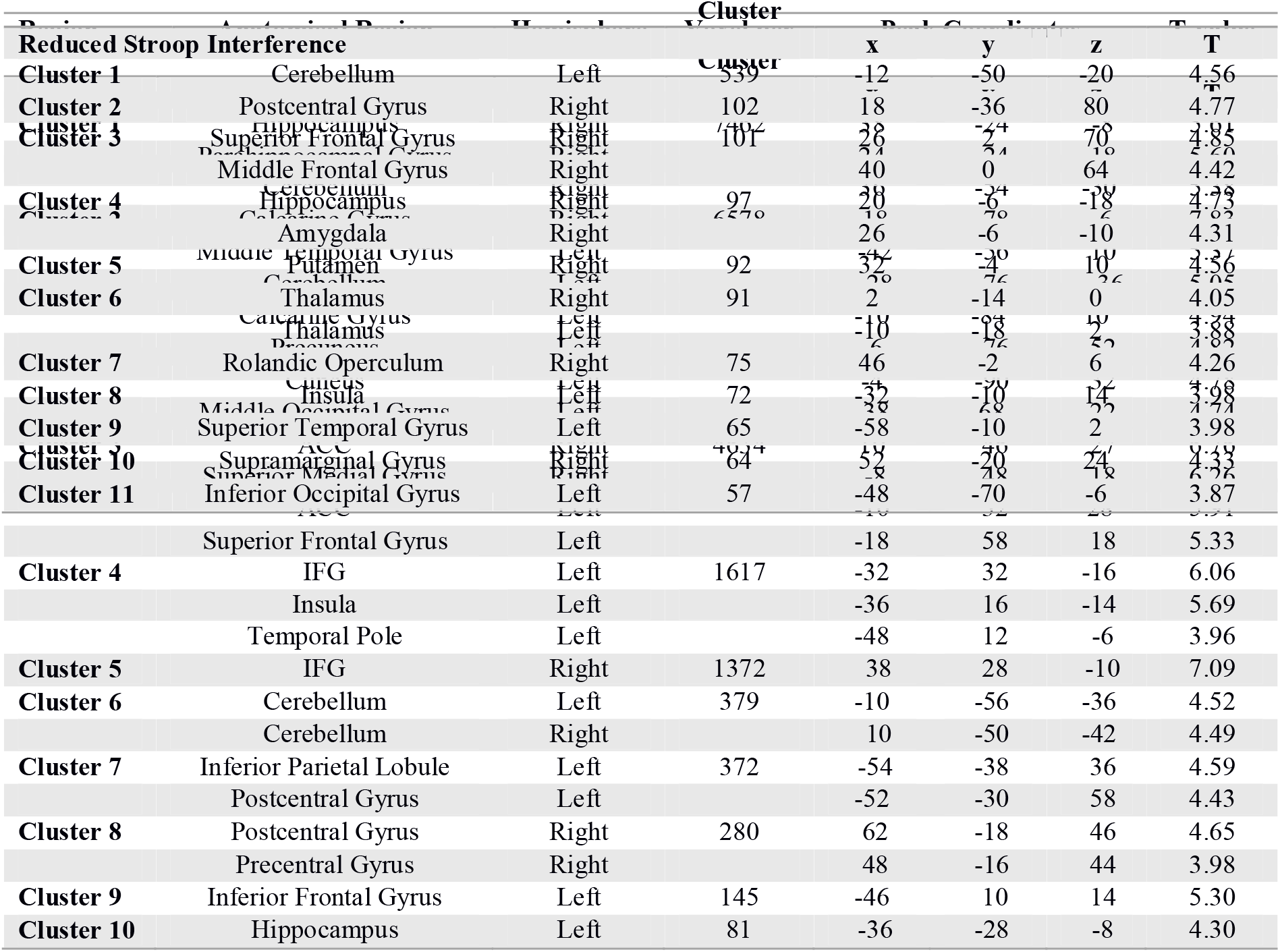
Clusterwise statistics of the Contrast FUN > TYP, including the clustersize, statistical thresholds, exemplary coordinates, T-value of the highest peaks and name of the activated area according to assigned regions – in the Anatomy toolbox

#### 3.2.2 Functional Activity: Covariate – Analysis (Stroop – Interference: FUN>TYP)

Analysis of covariance revealed a positive relationship between enhanced cognitive control abilities (low congruent – incongruent reaction times difference) and increased activity to humorous idea generation from the contrast FUN>TYP in several clusters, including the left hippocampus and the amygdala, the superior/medial frontal gyrus, the insula, the superior temporal gyrus, the IFG and the putamen (see Fig.3, Tab.2). No negative supra-threshold relationship was observed.

**Figure 3.**
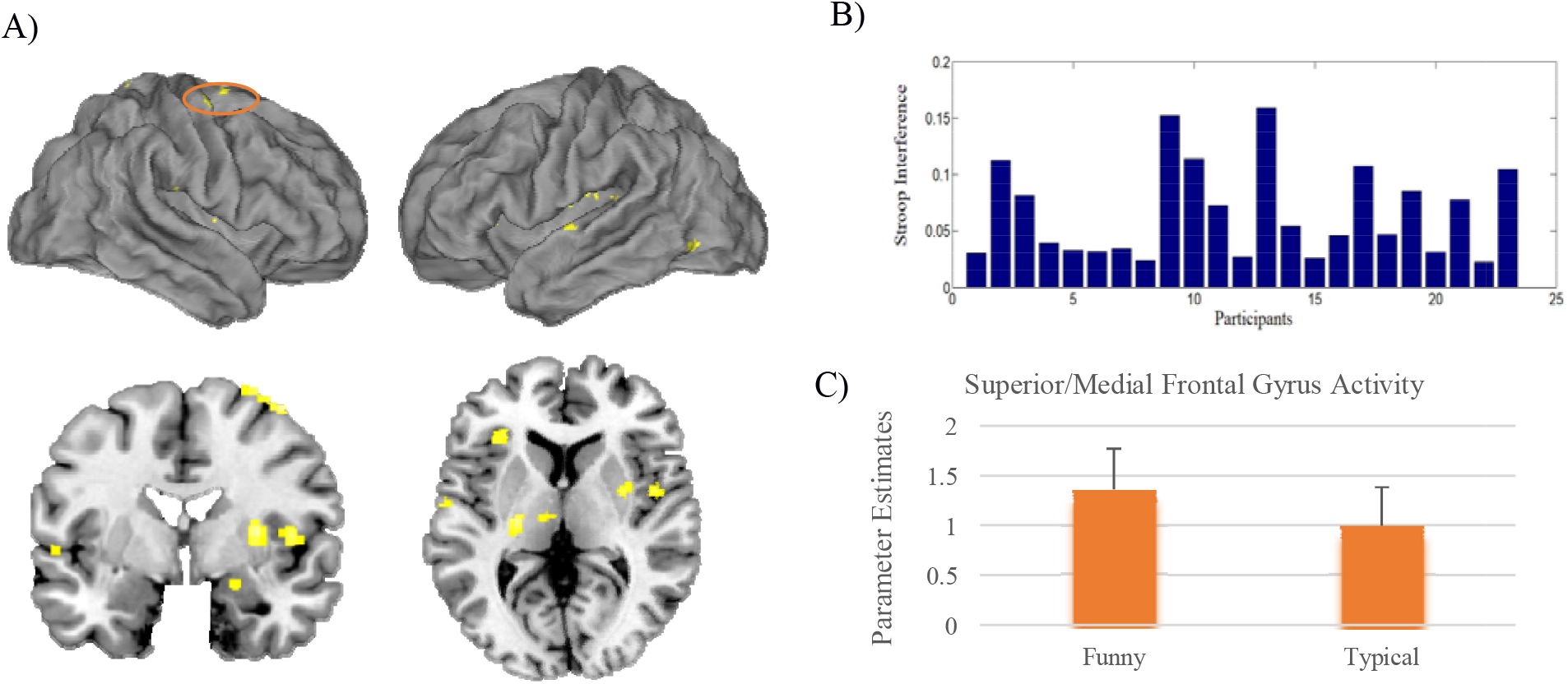
Functional Activity: Covariate Analysis (Stroop Interference): A) The functional covariation analysis of the stroop interference in the FUN>TYP contrast revealed a positive relationship between individual differences in cognitive inhibition and activation in several brain regions associated with humorous processing B) Distribution of the stroop interference values per participant. The value represents the difference between reaction times in the congruent vs. incongruent word-color pairs condition. C) Functional activity of the superior/medial frontal gyrus, the cluster was used for the subsequent connectivity analysis. Error bars represent SEM.

**Table 2.**
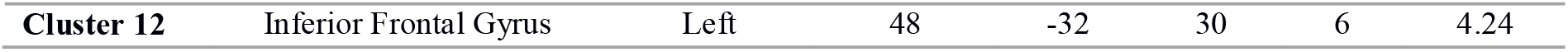
Clusterwise statistics of the analysis of covariance of the individual stroop score with the FUN > TYP contrast

**Table 3.** Clusterwise statistics of the functional connectivity analysis of covariance of the individual stroop score (reduced stroop interference) with the FUN>TYP contrast

| Region | Anatomical Region | Hemisphere | Voxel per Cluster | Peak Coordinates |  |  | T-value |
| --- | --- | --- | --- | --- | --- | --- | --- |
| Increased Stroop Interference |  |  | k | x | y | z | T |
| Cluster 1 | Precentral Gyrus | Right | 373 | 20 | -20 | 70 | 6.33 |
|  | Superior Frontal Gyrus | Right |  | 32 | -12 | 62 | 4.02 |
| Cluster 2 | Precentral Gyrus | Left | 371 | -48 | -2 | 34 | 6.03 |
|  | Postcentral Gyrus | Left |  | -46 | -18 | 30 | 4.32 |
| Small-Volume Correction | Left Amygdala | Left | 20 | -24 | -2 | -24 | 4.46 |

#### 3.2.3 Functional Connectivity: Covariate – Analysis (Stroop – Interference: FUN>TYP)

In a next step, the impact of cognitive control abilities on functional connectivity of the superior/medial frontal gyrus was assessed. This analysis revealed a reduced functional connectivity between the superior/medial frontal gyrus and the amygdala (small volume corrected, *t*=4.46, *p_FWE_*=.009) and the postcentral and precentral gyrus in participants with enhanced cognitive control abilities (Tab. 4), indicating a higher negative connectivity in individuals with increased cognitive control abilities.

#### 3.2.4 Amygdala Activity – Frontal Superior Medial Frontal Gyrus Connectivity Correlations

To investigate a potential increase of the amygdala activity by a negative functional coupling with prefrontal brain regions, we performed a correlation analysis between functional *activity* of the left amygdala and its functional *connectivity* with the superior/medial frontal gyrus. This analysis revealed a significant negative relationship between the amygdala activity and its connectivity with the superior/medial frontal gyrus (*r*=−.57, *p*<.05) indicating a stronger functional activity increase of the amygdala by a higher negative frontal coupling. The general relevance of this amygdala region for the generation of humorous ideas is shown by its enhanced activity in the funny compared to the typical condition (Region-of-Interest Analysis, *t*=1.73, *p*=.048, Fig. 5A).

**Figure 4.**
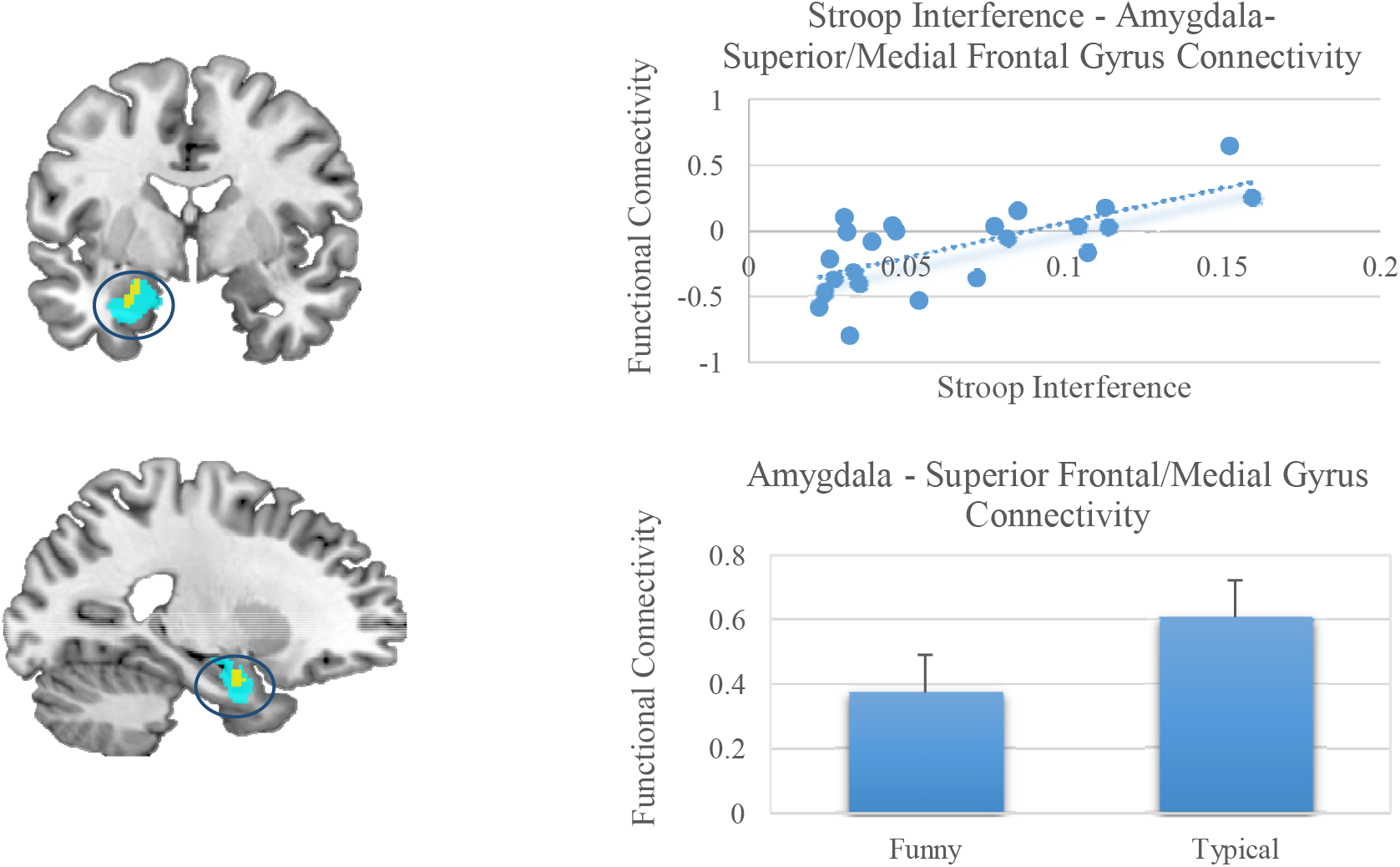
Functional Connectivity: Covariate Analysis (Stroop Interference): Significant functional connectivity between the superior frontal/medial gyrus and the amygdala (turquoise anatomically defined region-of-interest, yellow significantly activated amygdala (SMV-corrected)). Parameter Estimates were extracted from the significant functional amygdala cluster, the error bars represent SEM.

**Figure 5.**
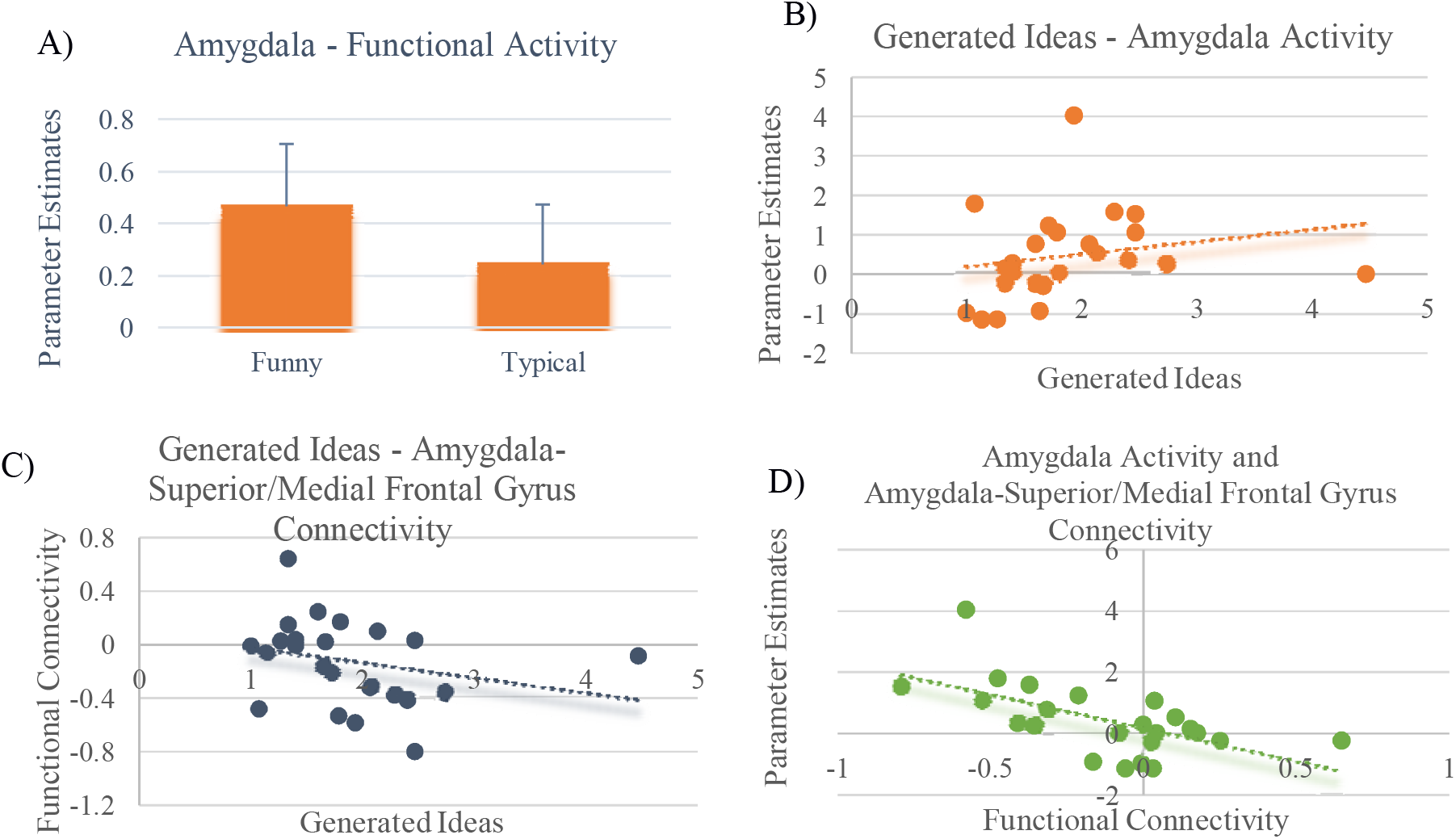
A) Significant higher functional activity in the amygdala in the funny vs. typical condition, the error bars represent SEM B) Higher amygdala activity was associated with increased generation of funny idea C) Reduced functional connectivity between the amygdala and the superior/medial frontal gyrus was associated with increased funny idea generation D) A higher negative functional connectivity between the amygdala – superior/medial frontal gyrus was associated with an increased functional activity in the amygdala.

#### 3.2.5 Brain – Behavior and Traits Correlations

Correlations of extracted beta estimates showed a significant association between the amygdala activity with the number of generated ideas in the funny condition (*r*=0.49, *p*<.05). Furthermore, the quantity of participants’ produced ideas during the funny condition correlated negatively with functional connectivity measures of the amygdala-superior/medial frontal gyrus (*r*=−0.44, *p*<.05), pointing to an increased productivity by a reduced functional connectivity. Additionally, we examined a potential association between the activity of the amygdala and behavioral tendencies to regulate emotions in real-life. This analysis showed that a suppressive emotion-regulation tendency (ERQ) (r=−.44, p<.05) is associated with reduced amygdala activity during the production of funny ideas.

## Discussion

In employing a novel humorous idea generation task and a classic stroop task, we identified the neural mechanisms underlying the production of funny ideas. Our results reveal that humorous idea production is associated with a brain circuit involving the amygdala, the hippocampus, the superior/medial frontal gyrus, the insula and the IFG. Remarkably, enhanced cognitive control abilities were associated with increased neural responses in the similar brain regions during the humor generation process, pointing to a facilitative effect of cognitive control abilities and abstract thinking. More specifically, we were able to demonstrate that participants with enhanced cognitive control abilities also displayed stronger negative functional connectivity between brain regions related to inhibitory processes (e.g. the superior/medial frontal gyrus; Hu et al., 2016) and the amygdala, a key region for humor processing (Berger et al., 2018b; Vritcka et al., 2013). Notably, the stronger the negative connectivity between the amygdala and brain regions related to cognitive control (i.e. superior/medial frontal gyrus) was, the greater the activity in the amygdala increased during funny idea generation. This finding is suggestive of an antagonism between brain regions related to cognitive control and the amygdala, leading to an increased availability of funny ideas. This interpretation is motivated by the finding that amygdala activity was linked both with outcome measures of the idea generation process, since participants with higher amygdala responses produced a higher amount of funny ideas and a tendency to use suppressive emotion-regulation strategies as a coping mechanism in real-life. Notably, our findings suggest that the well-established association between neural pathways linking cognitive control and affective systems (Ochsner et al., 2012), which has been found to be associated with increased negative emotional responses (Gold et al., 2014), is also relevant for the production of positive affective cognition and may therefore be particularly relevant for mental health (Martin, 1996).

Studies examining the cognitive and neural mechanisms of divergent thinking show that cognitive control mechanisms can facilitate the mining of abstract ideas (Jung et al., 2013). However, it has been speculated whether high (e.g. Edl et al., 2014) or low inhibitory mechanisms (Radel et al., 2015) enable the creative process (Benedek & Fink, 2019) and how these are represented in the brain. The assumption of a positive association between cognitive control processes and enhanced creativity (Benedek et al., 2014; Groborz & Necky, 2003) was supported by empirical findings showing that individuals with elaborated creative abilities also possess high cognitive control abilities in a stroop task (Edl et al., 2014). Moreover, a positive association between fluid intelligence and divergent thinking has been reported and linked to a higher creative potential (Benedek et al., 2014). The findings led to the theoretical interpretation that creative individuals are better able to suppress competing and obvious semantic associations, which is assumed to increase the availability of abstract concepts in the consciousness stream (Edl et al., 2014).

Adding to this picture, our imaging data indicate that participants with enhanced cognitive control abilities showed an increased amygdala activity but a negative connectivity with the superior frontal/medial gyrus, while generating humorous ideas. This finding suggests that an activity increase in the amygdala is associated with a superior frontal/medial gyrus activity decrease, and vice versa. This antagonistic *connectivity* was linked with a higher *activity* in the amygdala and related to an increase in idea generation. Our results are therefore compatible with findings from studies with psychiatric patients indicating that if cognitive and behavioral disinhibition is a core characteristics of the disorder, there is also a tendency for enhanced creative potential (Carson, 2011; Dietrich, 2014; Fink et al., 2014). A unifying element of disorders such as bipolar disorders (Green et al., 2007; Hajek et al., 2013), ADHD (Depue et al., 2010; Morein-Zamir, 2014) and schizophrenia (Hughes et al, 2012; Zandbelt et al., 2011) are functional and structural alterations in frontal brain regions associated with cognitive control processes. Taking these findings together, the studies suggest that an imbalance in inhibitory and disinhibitory processes is linked with a hypersensitivity in limbic structures (Clark & Sahakian, 2008; Plichta et al., 2009; Tan et al., 2012), which in turn is assumed to be associated with higher creativity (Carson, 2011; Dietrich, 2014; Flaherty, 2005). Interestingly, our data also suggests that a stronger amygdala response during generation of emotionally positive ideas is associated with an increased real-life usage of positive emotion-regulation strategies, i.e. reduced suppression of emotional responses which has been shown to be related to beneficial mental health outcomes (Gross & John, 2003). This association should therefore be further examined in psychiatric patients as, for example, interventions promoting humorous coping (Falkenberg et al., 2013) may to facilitate positive emotions in this group.

From a conceptual perspective, it has been suggested that frontal brain regions might facilitate the re-structuring of memory processes related to creative cognition (Benedek & Fink, 2019). This notion is somewhat contradictory to our finding of reduced frontal-amygdala connectivity being associated with an increased availability of funny ideas. Previous studies have pointed to a strong anatomical interconnectivity between the amygdala and the frontal cortex (Kim et al., 2011; Zimmermann et al., 2019), suggesting that frontal brain regions are implicated in the functional regulation of negative emotional and cognitive experiences, such as anxiety, triggered by the amygdala (Townsend et al., 2013; Li et al., 2018). Accordingly, our data indicate that in the domain of generating positive ideas a similar frontal downregulation is associated with an amygdala upregulation, and vice versa, which leads to an increase in the availability of positive ideas. Our interpretation is strengthened by an increased positive coupling of the amygdala with frontal brain regions during the control condition in which familiar and non-emotional concepts had to be retrieved. This suggests that in general inhibitory mechanisms of the amygdala control the expression of emotional associations (LeDoux, 2000; Quirk & Gehlert, 2003). Given that the amygdala is reliably found in humor processing (Vrticka et al., 2013) and considered a target of inhibitory mechanisms to regulate emotional states (Ehrlich et al., 2009) and memory processes (Yonelinas & Ritchey, 2015), we suggest that the regions key relevance during the generation of humor evolves by a dynamic coupling with brain regions relevant for inhibitory processes. Along this line, our findings open-up the question whether similar mechanisms can be found in clinical populations with schizophrenia, bipolar disorder or ADHD (Fink et al., 2014). With this study we provide the neurobiological key regions for the examination of related processes in clinical populations which might be extended by collecting the participants’ immediate experienced emotional state after generating humorous ideas in future studies.

## Acknowledgments

This work was supported by the Core Facility Brain Imaging, Faculty of Medicine, University of Marburg, Else Kröner-Fresenius-Stiftung (grant number: 2014_A136) and Deutsche Forschungsgemeinschaft (grant numbers: STR-1146/4-1, STR-1146/8-1, STR-1146/9-1, to BS)

## Competing interests

The authors report no competing interests.

## Notes

### Competing Interest Statement

The authors have declared no competing interest.

